# Isothermal Amplification of Nucleic Acids with Ladder-shape Melting Curve

**DOI:** 10.1101/2020.12.17.423190

**Authors:** Deguo Wang, Yongzhen Wang, Meng Zhang, Yongqing Zhang, Juntao Sun, Chumei Song, Fugang Xiao, Yuan Ping, Chen Pan, Yushan Hu, Chaoqun Wang, Yanhong Liu

## Abstract

A novel method termed isothermal am<u>p</u>lification of nucleic acids with <u>l</u>adder-shape <u>m</u>elting <u>c</u>urve (LMCP) was developed in the study. In this method, one pair of primers or two pairs of nested primers and a thermostable DNA polymerase (large fragment) were employed to amplify the <u>i</u>nternal <u>t</u>ranscribed <u>s</u>pacer (ITS) of *Oryza sativa* with ladder-shape melting curve. Our results demonstrated that the LMCP assay with nested primers was 50-fold more sensitive and one-hour faster than the LAMP assay with the same level of specificity. The LMCP method has the potential to be used for the prevention and control of the emerging epidemics.

## 1. Introduction

Pathogenic microorganisms are threats to the health of human and animals. For example, African Swine Fever Virus (ASFV) has brought great economical damage in swine industry, and 2019-nCoV has been leading to a global disaster with the death of millions people. The threat of ASFV, 2019-nCoV and newly emerging pathogenic microorganisms is emphatically still on, and a rapid, sensitive and simple nucleic acids test method is urgently needed.

Polymerase Chain Reaction (PCR) is the most commonly used method [1,2], e.g. for the nucleic acid detection of 2019-nCoV, but the application of this technology is limited due to the need of a thermal cycler. Isothermal amplification of nucleic acids does not need a thermal cycler, and some isothermal amplification methods have been elaborately designed, including nucleic acid sequence-based amplification (NASBA) [3], self-sustained sequence replication (3SR) [4], helicase-dependent amplification (HDA) [5], exponential amplification reaction (EXPAR) [6], Strand displacement amplification (SDA) [7], recombinase polymerase amplification (RPA) [8], rolling circle amplification (RCA) [9], and loop-mediated isothermal amplification (LAMP) [10], however, these methods have their own limitations including lengthy time and non-specific amplification [11,12]. Therefore, the nucleic acid test methods based on PCR are still dominant in the detection of 2019-nCoV epidemic, and great efforts have been made to modify the existing methods, e.g. enhanced recombinase polymerase amplification (eRPA) [13] and CRISPR–Cas9-triggered triggered SDA reactions [14].

In this paper, a novel method, termed isothermal amplification of nucleic acids with ladder-shape melting curve (LMCP), was developed. The reaction mechanism, target sequence selection, primer design, sensitivity and specificity of the amplification method are discussed. This new method can robustly amplify nucleic acids with high specificity and sensitivity in a very short time. Since this method does not need a thermocycler, it may have wide application for the detection of pandemic diseases including coronavirus.

## 2. Materials and Methods

### 2.1. Selection of the target sequences for nucleic acid detection

The nucleic acid sequences of *Oryza sativa* internal transcribed spacer (ITS) (GenBank ID: MF029734.1) were analyzed using Oligo 7 software (Molecular Biology Insights, Inc. Colorado Springs, CO, USA), and the target sequences with ladder-type melting curve were selected, as shown in Figure 1.

**Figure 1.**
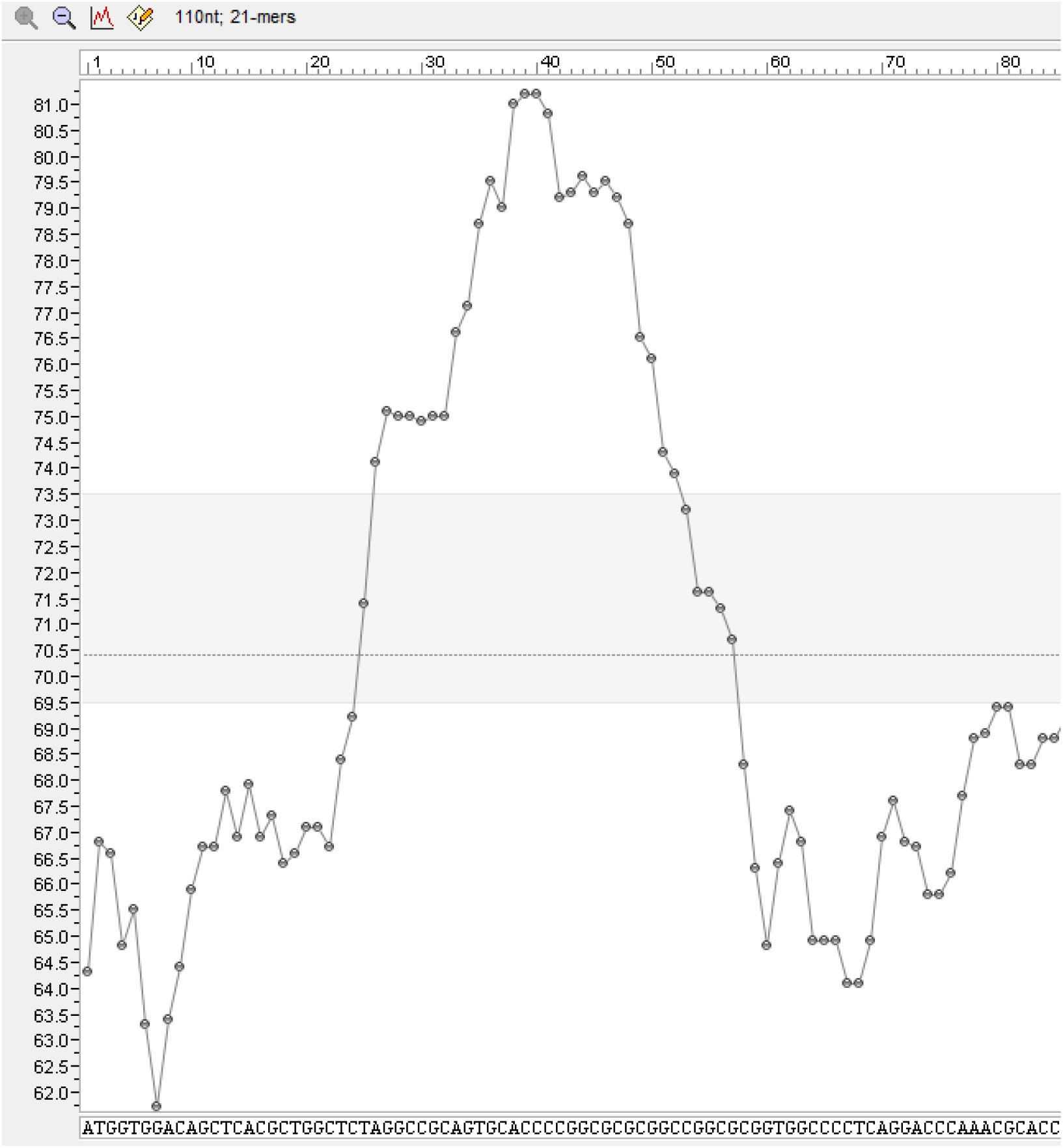
Melting Curve of Target Sequence (ITS) for Detection of *Oryza sativa* with LMCP Method.

### 2.2. Primer design for LMCP Assays

With the selected target sequences, a set of primers (forward primer and reverse primer, for short, F and R), No.1, was designed using the online software Primer3Plus (http://www.primer3plus.com/cgi-bin/dev/primer3plus.cgi); two pairs of nested primers (outer pair of primer P1 and D1, inner pair of primers P2 and D2) were designed using the same software, among which primer P1 and primer D2 were complementary to the minus strand while primer P2 and primer D1 were complementary to plus strand, the primer P (primer P2 and primer P1 linked with TTTT) and the primer D (primer D2 and primer D1 linked with -tttt-) were the No.2 set of primers, as shown in Table 1.

**Table 1.**
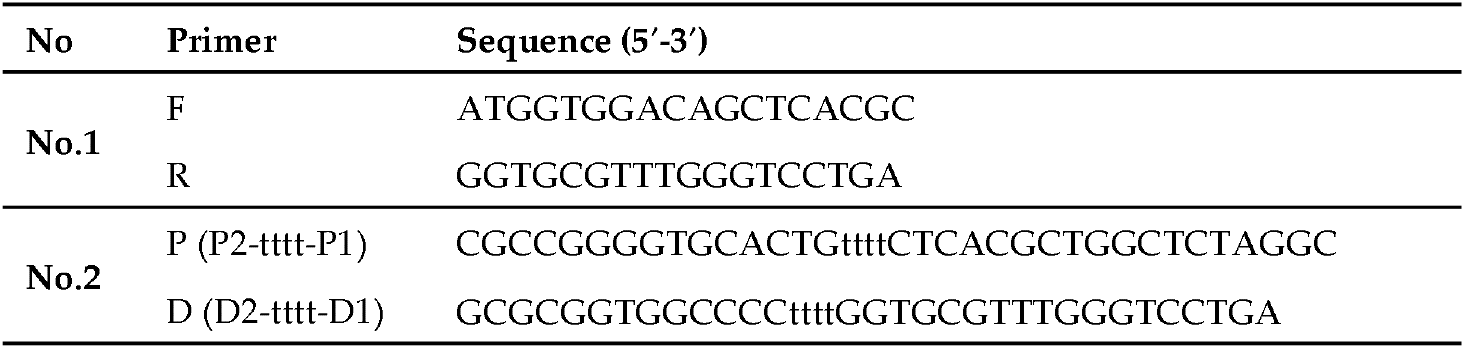
LMCP Primers for the ITS of *Oryza sativa*.

### 2.3. Genomic DNA isolation

Genomic DNA was isolated from *Oryza sativa*, *Zea mays*, *Vigna radiate*, *Glycine max*, *Ipomoea batatas*, *Solanum tuberosum* and *Manihot esculenta* using NucleoSpin® Food kit (MACHEREY-NAGEL GmbH & Co. KG) according to the manufacturer’s instruction.

### 2.4. Sensitivity comparison of the LMCP and LAMP assays

The LMCP assays with different sets of primers were performed in a 10-μL reaction mixture containing 0.8 mM of each primers (No.1 set of Primer F and Primer R, or No.2 set of Primer P and Primer D), 1.0 mM dNTPs, 1× reaction buffer (20 mM Tris-HCl (pH 8.8), 10 mM KCl, 10 mM (NH_4_)_2_SO_4_, 6 mM MgSO_4_, 0.1% Triton X-100) and 3.2 U Bst DNA polymerase (Merit Biotech (Shandong) Co., Ltd, China), and 1×SYBR Green I. The detection limits of LMCP assays with No.1 or No.2 sets of primers were determined with 10-fold dilution of genomic DNA from *Oryza sativa*, the reaction mixtures with No.1 set of primers were heated at the optimized temperature 56.5°C for 100 min (2 min per cycle) in StepOne™ System Real-Time PCR System (Applied Biosystems, Foster City, CA, USA), while the reaction mixtures with No.2 set of primers were heated at 55°C for 60 min (30 s per cycle), respectively. A melting curve was obtained for each reaction.

The LAMP primers targeting the *Oryza sativa* internal transcribed spacer (ITS) (GenBank ID: MF029734.1) were designed using PrimerExplorer V5 (http://primerexplorer.jp/e/) (Please see Table 2 for primer sequences). The LAMP assay with different set of LAMP primers was performed in a 10-μL reaction mixture containing 0.8 mM each of forward inner primer (FIP) and backward inner primer (BIP), 0.2 mM each of forward outer primer (F3) and backward outer primer (B3), 1.0 mM dNTPs, 1× reaction buffer (20 mM Tris-HCl (pH 8.8), 10 mM KCl, 10 mM (NH_4_)_2_SO_4_, 6 mM MgSO_4_, 0.1% Triton X-100) and 3.2 U Bst DNA polymerase (Merit Biotech (Shandong) Co., Ltd, China), and 1×SYBR Green I. The detection limit of LAMP assay was determined with 10-fold dilution of genomic DNA from *Oryza sativa*, and the reaction mixture was heated at the optimized temperature 59°C for 2 h (90 s per cycle).A melting curve was obtained using a StepOne™ System (Applied Biosystems, Foster City, CA, USA).

**Table 2.**
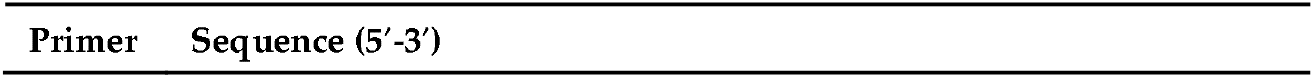
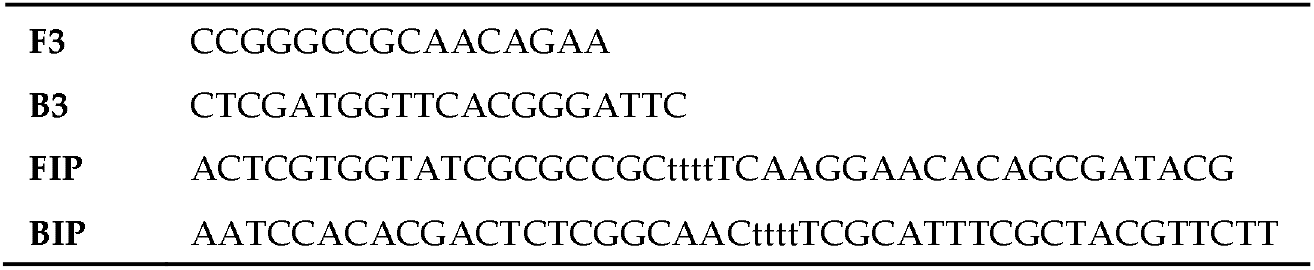
LAMP Primers for the ITS of *Oryza sativa*.

### 2.5. Specificity Comparison of the LMCP and LAMP Assays

Genomic DNA from *Oryza sativa*, *Zea mays*, *Vigna radiate*, *Glycine max*, *Ipomoea batatas*, *Solanum tuberosum and Manihot esculenta* were randomly selected for the specificity determination of the LMCP assay with No.1 set of primers, No.2 set of primers and the LAMP assay, and the amount of genomic DNA used was 1 ng per reaction.

## 3. Results

### 3.1. Principle of the LMCP method

The major mechanism, also the greatest progress compared with the previously developed nucleic acid amplification methods, was the non-thermal and non-enzymatic single-strand template supply. The nucleic acid sequences with ladder-type melting curve were selected as target, as shown in Figure 2, the Tms of P1 and/or D1 were/was higher than the amplification temperature, whereas the Tms of corresponding fragments/fragment of P1 and/or D1 on the target were/was lower than the amplification temperature, and the single-strand template was supplied via the Tm difference.

**Figure 2.**
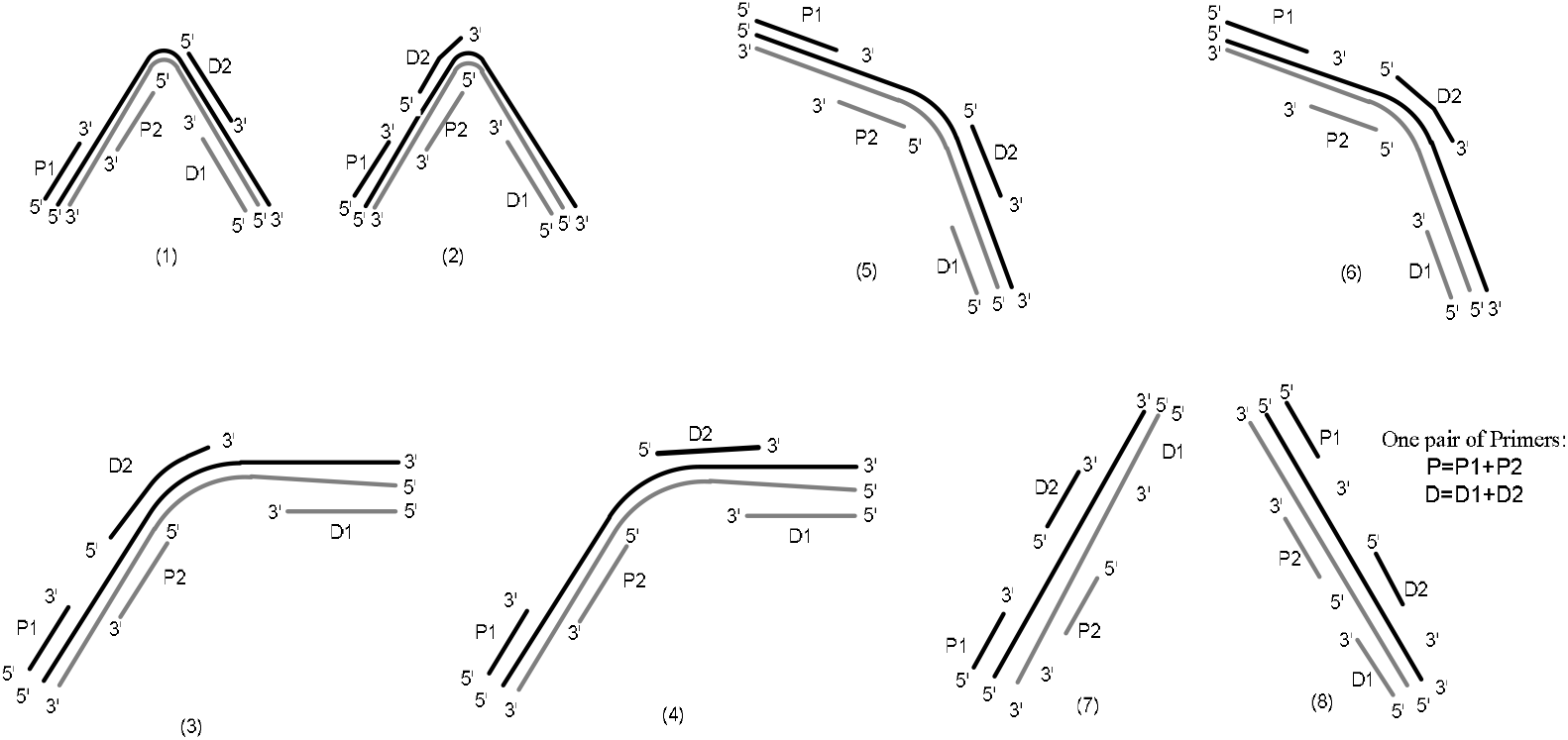
The Ladder-type Melting Curves of LMCP and their Primers.

When the Tm of both P1 and D1 were higher than the amplification temperature and the Tm of both corresponding fragments were lower than the amplification temperature, the two primers can amplify the target like PCR without temperature cycles of denaturation, annealing and extension.

For all of the ladder-type nucleic acid targets in Figure 2, the target sequences can be amplified by the pair of primer P (primer P2 and primer P1 linked with TTTT) and primer D (primer D2 and primer D1 linked with -tttt-). The whole amplification process can be divided into initial and exponential amplification stages. In the initial stage, when only the Tm of P1 was higher than the amplification temperature while the Tm of corresponding fragment of P1 on the target was lower than the amplification temperature, as shown in Figure 3, the P1 on the primer P annealed with the complementary fragment via the Tm difference initiated synthesis reaction and strand displacement reaction with the action of thermostable DNA polymerase (large fragment), then the P1 on the primer P annealed with the complementary fragment of newly synthesized double strand DNA and initiated new cycle of synthesis reaction and strand displacement reaction, while the D1 of primer D annealed with the displaced single strand DNA and synthesized a double-strand DNA, after a series of annealing, synthesis and strand displacement, the initial structure for the exponential amplification stage was formed, which was a dumb-bell form of DNA. When the Tms of both P1 of primer P and D1 of primer D were higher than amplification temperature while the Tms of their corresponding fragments were lower than the amplification temperature, the initial stage produced the dumb-bell form DNA for the exponential amplification stage as above, as shown in Figure 4.

**Figure 3.**
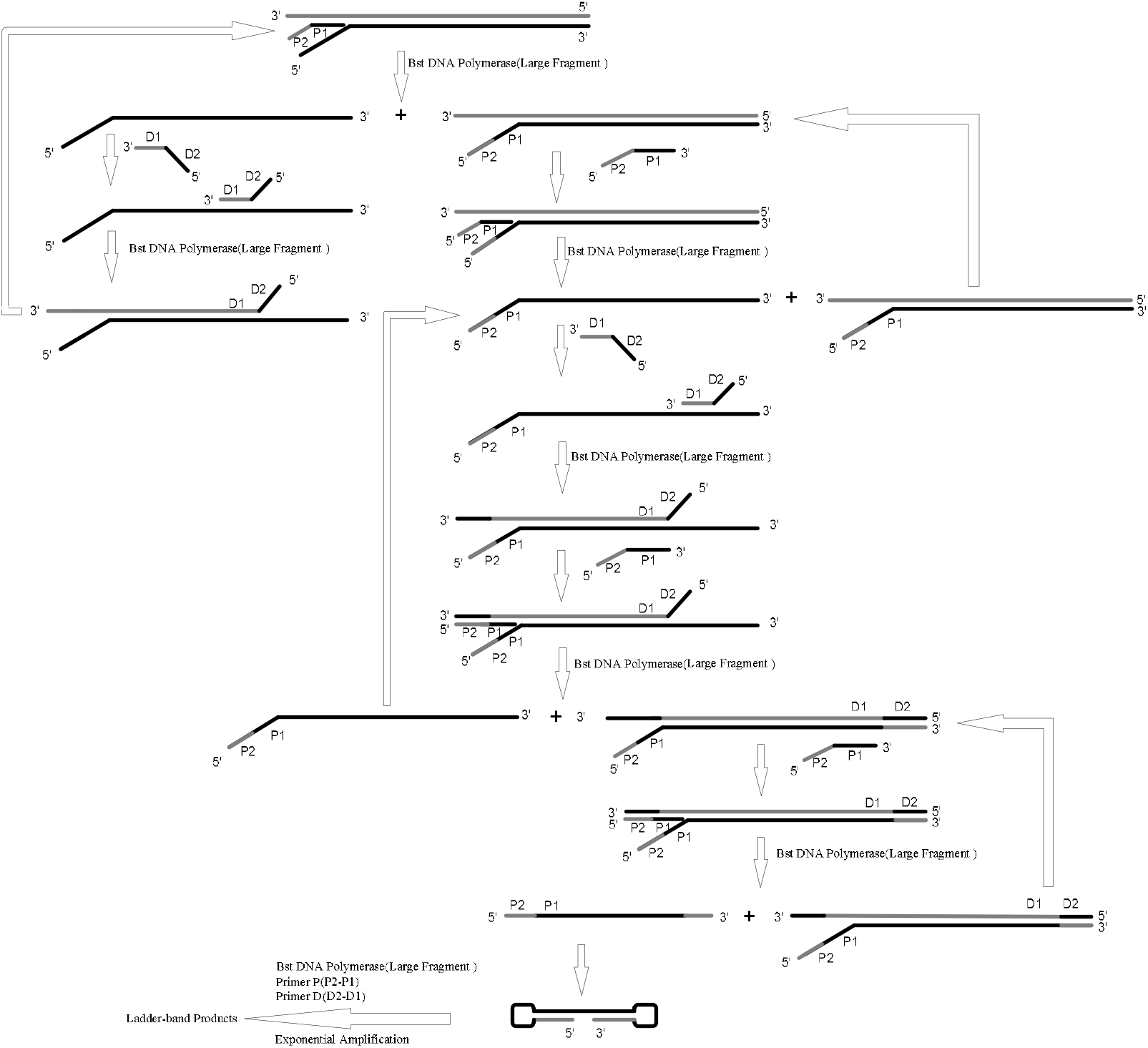
Schematic representation (1) for Initial Stage of LMCP. Note: The Tm of P1 was higher than the amplification temperature and the Tm of corresponding fragment of P1 was lower than the amplification temperature at the same time.

**Figure 4.**
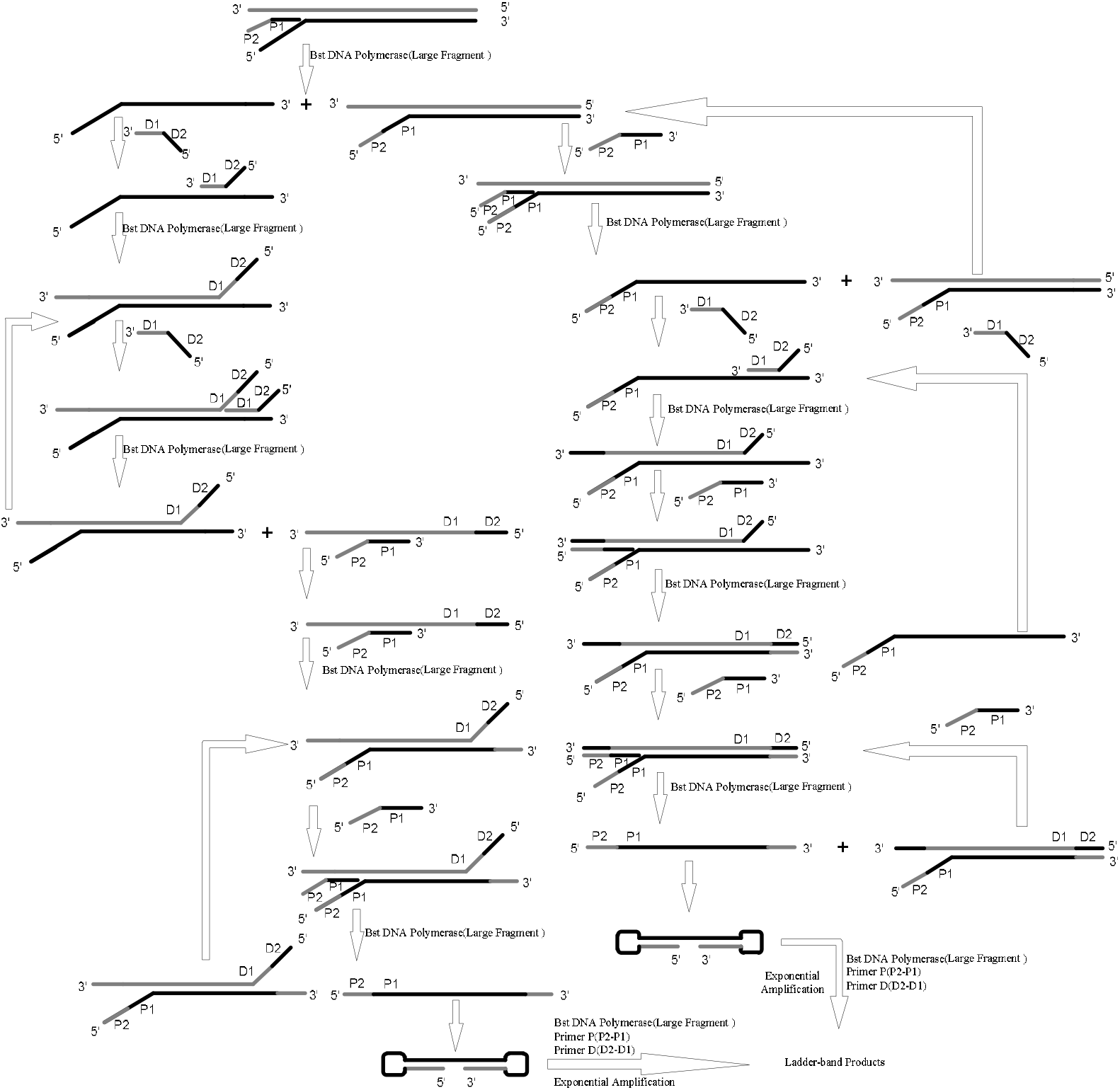
Schematic representation (2) for Initial Stage of LMCP. Note: The Tms of P1 and D1 were higher than the amplification temperature whereas the Tms of corresponding fragments of P1 and D1 were lower than the amplification temperature.

In the exponential amplification stage (Figure 5), the 3’ end of dumb-bell form of DNA initiated self-primed synthesis and strand displacement reaction with the action of thermostable DNA polymerase (large fragment), and produced the double strand DNA with a blunt end and a single-strand stem-loop end (Step 1); the D1 of primer D annealing with the stem loop of the newly formed double-strand DNA via complementary base pairing initiated synthesis reaction and strand displacement reaction with the action of thermostable DNA polymerase (large fragment) (Step 2), and produced the semi-single-strand and semi-double-strand DNA with a blunt end, middle double-strand stem loop and a single-strand stem-loop end (Step 3); the 3’ end of newly formed structure initiated self-primed synthesis and strand displacement reaction with action of thermostable DNA polymerase (large fragment) (Step 4), and produced a double-strand DNA with a blunt end, middle double-strand stem loop and a single-strand stem-loop end and a single-strand DNA, P1 of primer P annealing with the single-strand stem loop of newly produced double-strand DNA initiated a cycle of synthesis reaction and strand displacement reaction (Step 5, Step 6, Step 7…), and the newly formed single strand DNA quickly converted to dumb-bell form DNA via two-end complementary base pairing and initiated a new cycle of self-primed synthesis and strand displacement reaction with action of thermostable DNA polymerase (large fragment) (Step 9, Step 10…); and the numerous DNA with length as different-time as that of target sequence produced after a series of self-primed synthesis, annealing, synthesis and strand displacement reactions.

**Figure 5.**
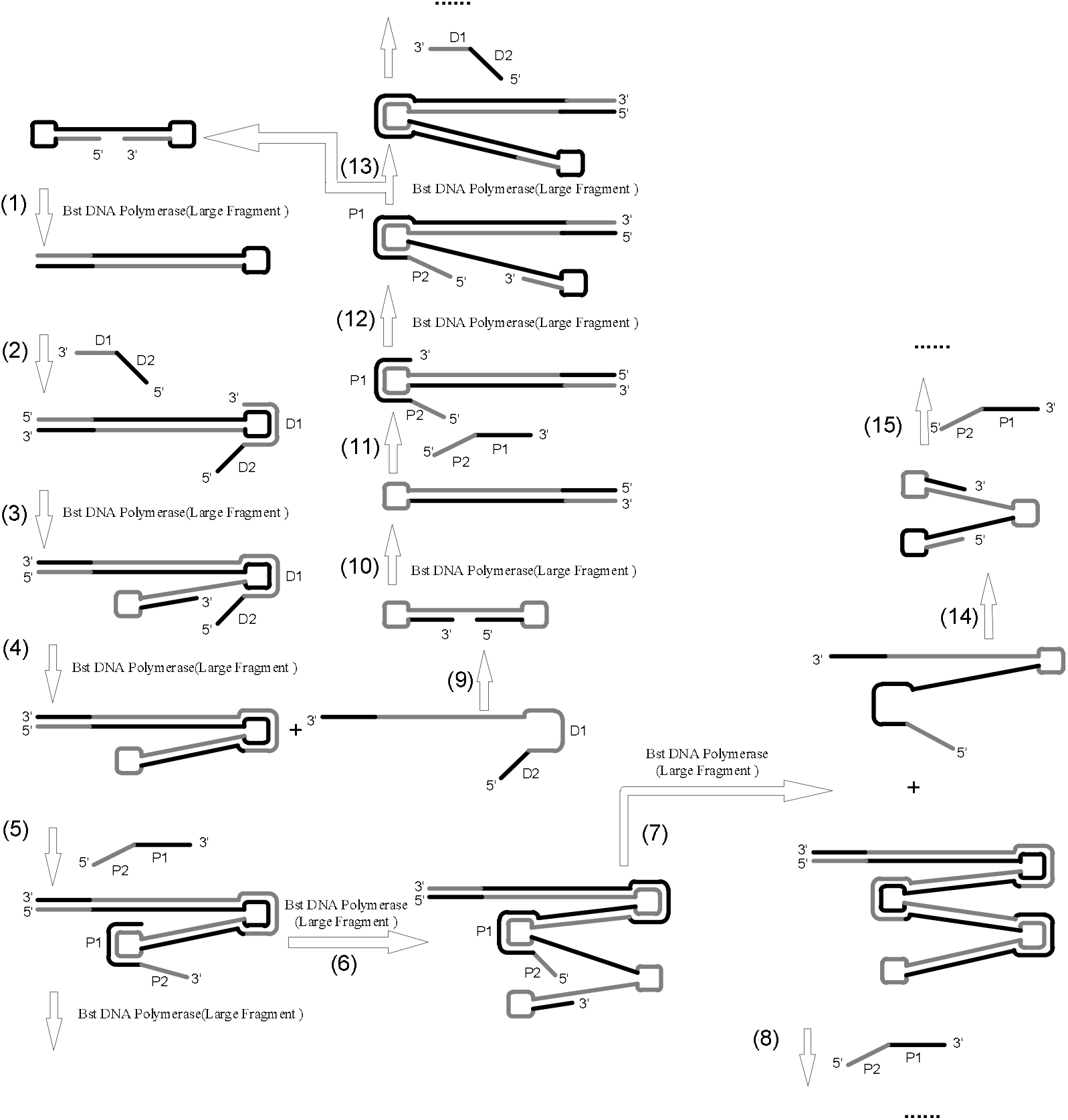
The Diagrammatic Sketch (2) for Exponential Amplification Stage of LMCP.

### 3.2. Sensitivity of the LMCP and the LAMP assays

The detection limits of the LMCP assays with No.1 set of primers and No.2 set of primers as well as the LAMP assay were determined using 10-fold dilution of genomic DNA of *Oryza sativa*. In the case of non-specific amplification judged via the melting curves, the real detection limits of the LMCP with No.1 set of primers and No.2 set of primers as well as the LAMP assay were 100 pg, 10 pg and 500 pg genomic DNA of *Oryza sativa* (Figure 6 and Figure 7), the LMCP assay with No.2 set of primers was 50-fold more sensitive than the LAMP assay, and one-hour more rapid than the LAMP assay.

**Figure 6.**
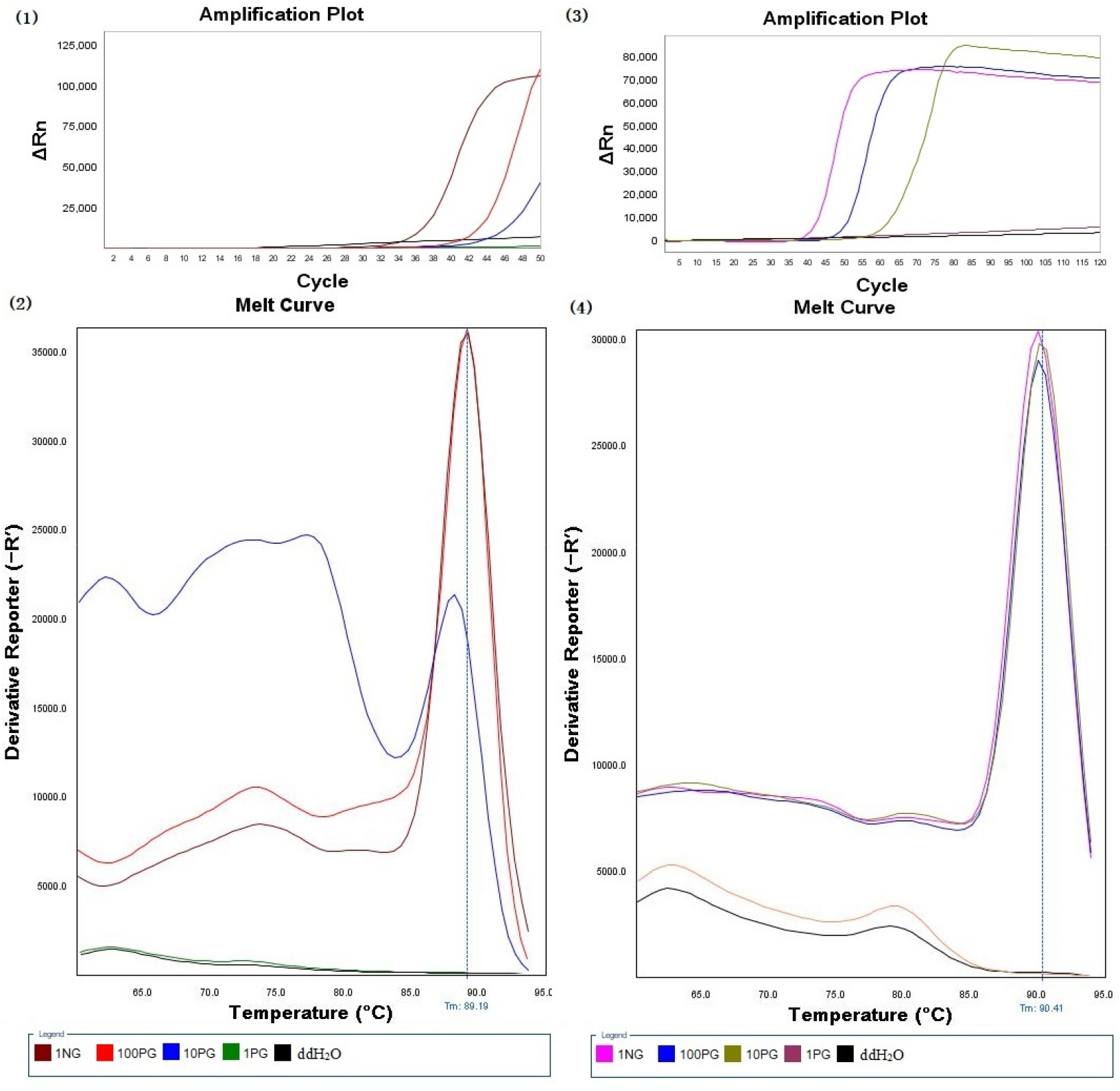
Sensitivity determination of LMCP assays for detection of *Oryza sativa* ITS using StepOne™ System Real-Time PCR System. (1). Amplification plot of LMCP assay with No.1 set of primers; (2). Melting curve of LMCP assay with No.1 set of primers; (3). Amplification plot of LMCP assay with No.2 set of primers; (4). Melting curve of LMCP assay with No.2 set of primers.

**Figure 7.**
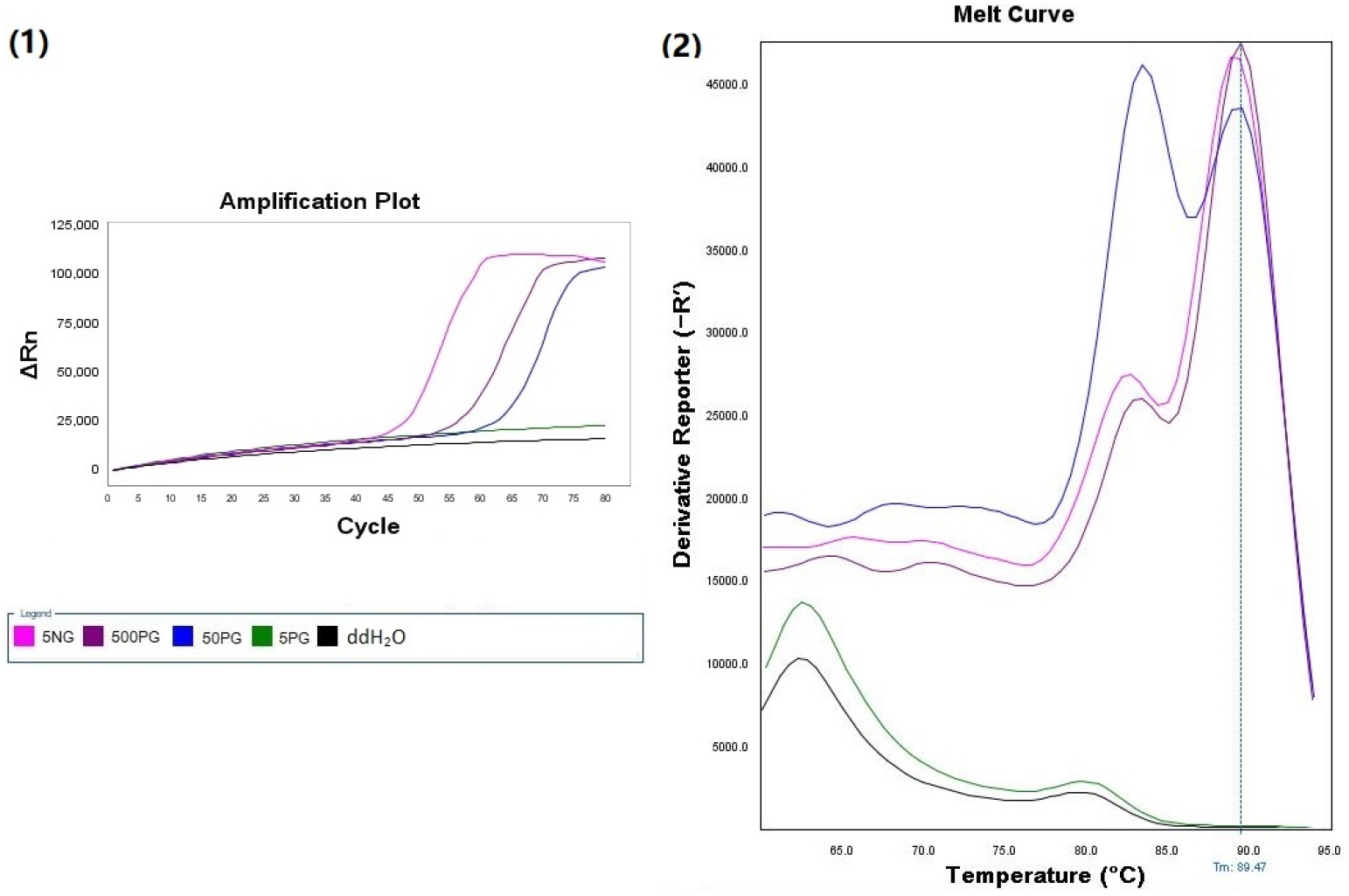
Sensitivity determination of LAMP assay for detection of *Oryza sativa* ITS using StepOne™ System Real-Time PCR System. (1). Amplification plot of LAMP assay; (2). Melting curve of LAMP assay with No.1 set of primers.

### 3.3. Specificity of the LMCP and LAMP assays

As shown in Figure 8 and in Figure 9, both the LMCP assays, as similar as the LAMP assay, were of high specificity, which only amplified the genomic DNA from *Oryza sativa*.

**Figure 8.**
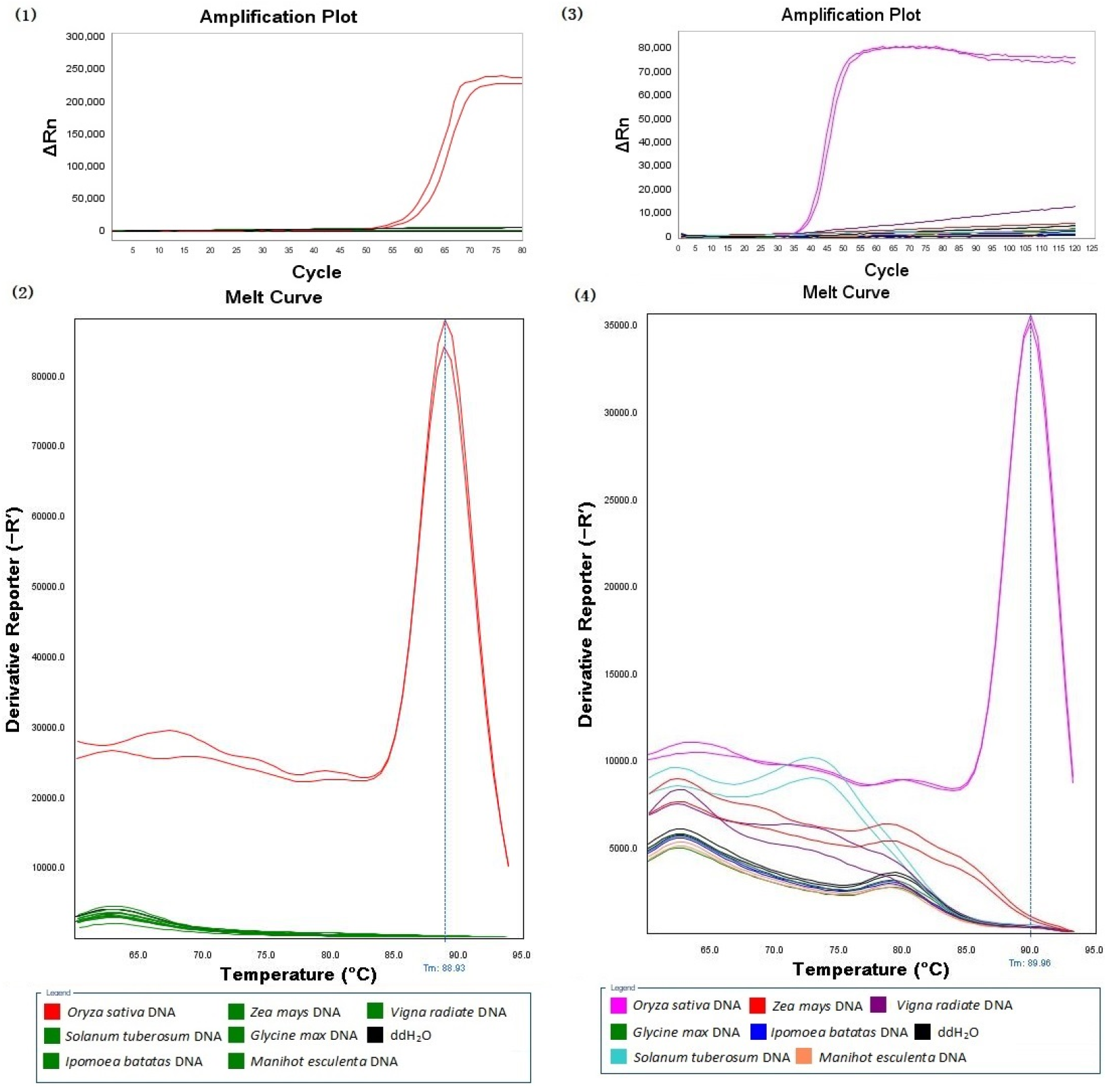
Specificity determination of LMCP assays for detection of *Oryza sativa* ITS using StepOne™ System Real-Time PCR System. (1). Amplification plot of LMCP assay with No.1 set of primers; (2). Melting curve of LMCP assay with No.1 set of primers; (3). Amplification plot of LMCP assay with No.2 set of primers; (4). Melting curve of LMCP assay with No.2 set of primers.

**Figure 9.**
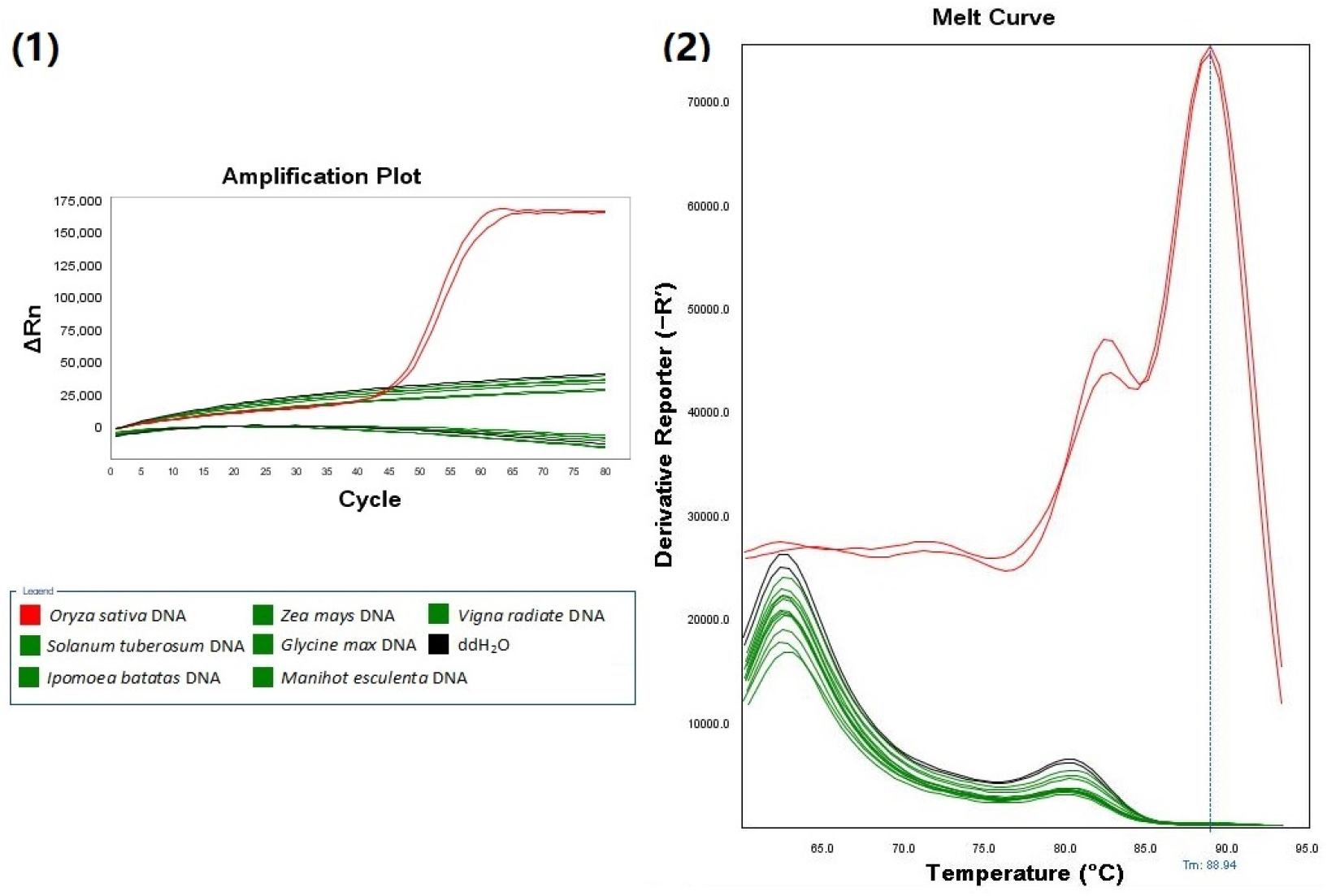
Specificity determination of LAMP assay for detection of *Oryza sativa* ITS using StepOne™ System Real-Time PCR System. (1). Amplification plot of LAMP assay; (2). Melting curve of LAMP assay.

## 4. Discussion

Compared with previously established detection methods, the LMCP developed in this study offers following advantages: **(i) Simple Mechanism**, this method neither needs the thermal denaturation in PCR [1,2] nor the enzymes other than thermostable DNA polymerase (large fragment), e.g. reverse transcriptase in Nuclear Acid Sequence-based Amplification [3,6], NEase in exponential strand displacement amplification (E-SDA), primer generation-rolling circle amplification (PG-RCA) and EXPAR [6,15, 16], ligase in ligation-mediated hyperbranched rolling circle amplification (HRCA) [17], helicase in HAD [5] and recombinase in RPA [8]. **(ii) Easy Primer Design**, the primers of LMCP can be easily designed with the commonly used PCR primer designing software. **(iii) Quick turn-around time**, the reaction time of LMCP with nested primers is less than half an hour; while the reaction time of most previous developed methods is more than 0.5 h [11], the reaction time of LAMP in the study was approximate 1.5 h. **(iv) High Specificity**, the specific primer sequences, special requirement for Tm of primer and its corresponding fragment and particular requirement for the melting curve shape of target sequence lead to the high specificity of the LMCP. **(v) High Sensitivity**, the detection limit of LMCP assays either with one pair of primers or with two pairs of nested primers is 3-5 copies per reaction. **(vi) Low Requirement for Target Sequence Length**, the target sequence length of LMCP is generally≥60nt, e.g. the target sequence of No.2 set of primers in this study is 76 nt, while most LAMP methods generally require the target sequence to be more than 200 nt [10]. **(vii) Wide Application Range**, the LMCP can take single-strand DNA, double-strand DNA or RNA as targets. It is easy to understand that the LMCP can take single-strand DNA or double-strand DNA as target, which has been clearly explained and experimentally confirmed in this study. The reason that the LMCP can amplify RNA is that the Bst DNA polymerase also possesses the activity of reverse transcriptase [18], which needs further study. The ITS of *Oryza sativa* was used to developed LMCP assay as a proof of concept in this study. Due to the low requirement of target sequence length and no requirement for a thermocycler, the LMCP assays can be extended to other emerging pandemics possibly 2019-nCoV.

## 5. Conclusions

The novel method, isothermal amplification of nucleic acids with ladder-shape melting curve (LMCP), was developed in this study, which was 50-fold more sensitive and one-hour faster than the LAMP assay with the same level of specificity.

## Author Contributions

D.-G.W. and Y.-Z.W. conceived the experiments. M.Z., Y.-Q.Z. and J.-T.S., established the setup and performed the experiment. Y.-S.H., Y.P. and P.C. analyzed the data and discussed the results. D.-G.W., Y.-Z.W., and Y.-H.L. wrote the manuscript. All authors read the manuscript. D.-G.W. and Y.-H.L. supervised the project.

## Funding

The research was funded by the National Key Research and Development Program of China, grant number 2016YFD0500704, and by Henan Science and Technology Plan Project, grant number 182102110285.

Informed Consent Statement: “Not applicable” for studies not involving humans.

## Data Availability Statement

The authors declare that all data generated or analyzed during this study are included in this published article and its Supplementary Information files. A Source Data file is available. Any other relevant data is available upon reasonable request.

## Acknowledgments

We thank all members from Key Laboratory of Biomarker Based Rapid-detection Technology for Food Safety of Henan Province, and would like to thank Prof. Zihe Rao insightful discussion.

## Conflicts of Interest

The authors declare that they have no competing interests.

## Notes

### Competing Interest Statement

The authors have declared no competing interest.

